# Processing and sectioning of organ donor spinal cord tissue for electrophysiology on acute human spinal cord slices

**DOI:** 10.1101/2025.07.26.666132

**Authors:** Annemarie Dedek, Eder Gambeta, Raksha Shriraam, Emine Topcu, Jeff S. McDermott, Jeffrey L. Krajewski, Eve C. Tsai, Michael E. Hildebrand

**Author notes:** Corresponding Author: Michael Hildebrand, PhD Professor, Department of Neuroscience, Carleton University Associate Vice-Provost (Graduate Student Affairs) Affiliate Investigator, Ottawa Hospital Research Institute 1125 Colonel By Drive 6310 Health Sciences Building Ottawa, ON K1S 5B6 CANADA.

## Abstract

Acute spinal cord slice electrophysiology is a powerful technique used in preclinical basic science research to investigate sensory and motor neuron function and pathophysiology. A major barrier that stands between implementing these findings into effective clinical treatments is the translational gap between rodent models and human patients. To date, no methods or protocols describe how to prepare viable human spinal cord slices for acute electrophysiological recordings. To bridge this translational divide, we describe here a protocol for the extraction of spinal cord tissue from consenting human organ donors and the preparation and sectioning of this tissue for acute spinal cord slice electrophysiology. With the collaboration of a transplant service and licensed surgeon, tissue can be extracted in 30-50 minutes. Acute spinal cord slices can then be prepared in the laboratory by trained graduate students in 2.5-5 hours, depending on the amount of tissue and scope of experiments. Using a viability stain to confirm that spinal slices are of sufficient quality to proceed, slices can then be used for either patch-clamp recordings to study the excitability of individual neurons or for high-density multielectrode array recordings to study intact sensory circuits. Slices remain viable for 4 to 8 hours, providing ample time for investigating synaptic and circuit-level signalling dynamics, including the use of pharmacological agents to probe the roles of specific molecular targets. The approaches described here can be implemented to improve translational physiological research and as a human tissue-based preclinical drug target identification and validation assay.

## Introduction

Studying the molecular determinants of sensory and motor processing in rodent preclinical models is a key first step towards the identification of new treatment approaches for diseases of the spinal cord. Preclinical human tissue assays are emerging to bridge the translational divide between these rodent models and target human populations, with a protocol for culturing human peripheral sensory neurons (Valtcheva et al., 2016) leading to transformational advances in understanding and targeting peripheral mechanisms of pain. Within a decade of using this published protocol (Valtcheva et al., 2016), researchers across the globe have identified species differences in: 1) sensory neuron subpopulations (Bhuiyan et al., 2024; Nguyen et al., 2021; Ray et al., 2018; Shiers et al., 2020; Tavares-Ferreira et al., 2022), 2) the upregulation of inflammation-associated genes (Wangzhou et al., 2020), and 3) the expression and function of ion channels and receptors (McIlvried et al., 2025; Moy et al., 2020; Rostock et al., 2018; Sheahan et al., 2018; Yi et al., 2023, 2025; Zurek, Thiyagarajan, et al., 2024). Furthermore, human sensory neuron assays are now being used to validate therapeutic targets and to test novel compounds. For example, Yi and colleagues uncovered that human sensory neurons from donors with a history of pain express higher levels of RNA transcripts encoding the voltage-gated sodium channel 1.8 (Nav1.8) (Yi et al., 2025). In parallel, testing candidate antagonists of Nav1.8-mediated currents in human sensory neurons led to the development of a novel human-specific Nav1.8 inhibitor, Suzetrigine, which was recently approved by the U.S. Federal Drug Administration for the treatment of moderate to severe acute pain (Copits et al., 2025; Jones et al., 2023). These peripheral neuron examples reinforce a shift in both fundamental science and industrial drug development towards more predictive human tissue preclinical assays (Renthal et al., 2021).

The spinal cord is a critical hub for sensory and motor processing (Abraira & Ginty, 2013; Bizzi et al., 2000; D’Mello & Dickenson, 2008; Fitzgerald & Jennings, 1999; Furue et al., 2004; Koch et al., 2018; Poppele & Bosco, 2003; Tuthill & Azim, 2018), with many molecular targets being identified (Attal et al., 2023; Bartlett et al., 2020; Bouhassira & Attal, 2018; C. Montana & W. Gereau, 2011; Li & Zhang, 2012; Marwaha et al., 2016; Obeng et al., 2021; Scolding et al., 2017; Wang et al., 2019), but little success in clinical translation to date (Bartlett et al., 2020; Eisenach & Rice, 2022; Khanna, 2012; Kiernan et al., 2020; Lubetzki et al., 2020; Oshinsky et al., 2022). Although most fundamental spinal cord research has relied on rodents, our pioneering use of human spinal cord (hSC) physiology assays (Davis et al., 2025; Dedek et al., 2019, 2022, 2024; Parnell et al., 2023; Yadav et al., 2023) has opened the door to addressing these critical translational questions, similar to that done for human peripheral sensory neurons. These studies have allowed us to compare receptor expression (Parnell et al., 2023) and neuronal biophysical properties (Dedek et al., 2024) across sex and species under non-pathological conditions, with many features being conserved across species and sex, but also some clinically-relevant differences in receptor expression and function between rodents and humans (Dedek et al., 2024). When characterizing signalling pathways in an *ex vivo* human tissue model of chronic pain, we uncovered that molecular determinants of pathophysiology are conserved across species (Dedek et al., 2019, 2022), but differ across sex (Dedek et al., 2022). We have also used our hSC assays to investigate cellular taxonomy across species (Yadav et al., 2023), as well as to compare the dorsal horn expression of a candidate therapeutic target, KCC2, between healthy donors and those with a history of pain or long-term opioid use (Davis et al., 2025).

Despite the explosion of human sensory neuron research (Bhuiyan et al., 2024; Davidson et al., 2014; Hartung et al., 2022; McIlvried et al., 2023, 2025; Moy et al., 2020; Ray et al., 2018; Sheahan et al., 2018; Shiers et al., 2020; Tavares-Ferreira et al., 2022; Valtcheva et al., 2016; Wangzhou et al., 2020; Yi et al., 2024; X. Zhang et al., 2017, 2019; X. L. Zhang et al., 2015; Zurek, Ehsanian, et al., 2024; Zurek, Thiyagarajan, et al., 2024), there is a lack of physiology studies using hSC tissue. A particularly challenging technique to incorporate into hSC assays is electrophysiology, which is widely used in both physiological and pharmacological studies to examine synaptic transmission, ion channel activity, membrane potential changes, and network activity. These functional outputs also provide opportunities to identify druggable targets and to test the efficacy of candidate drugs in human preclinical assays. Unlike the human sensory neurons described above, physiological studies on human CNS tissue have been mainly restricted to extracted pathological tissue in patients with epilepsy or brain tumors (Chameh et al., 2023; Howard et al., 2022; Kushner et al., 2022) and are limited by the typically compromised cellular viability within the tissue that is collected. Here, we detail a protocol that can be used to prepare high-quality acute hSC slices collected from organ donors for either patch-clamp or high-density multi-electrode array (hdMEA) electrophysiological recordings on healthy dorsal horn neurons. In addition, tissue that is not used for electrophysiology can be flash-frozen or fixed for use in parallel genetic, biochemical, or immunohistochemical experiments.

## Development of the protocol

Building off of previous work on human sensory neurons (Valtcheva et al., 2016) and a protocol for culturing neural stem/progenitor cells (NSPCs) from the central canal of the human spinal cord (Galuta et al., 2020), we have developed an assay for the preparation of viable hSC slices for electrophysiological recording. Similar to previous studies, this protocol employs an anterior surgical extraction approach, which provides the benefit of not requiring any additional incisions to be made to the skin of the organ donor, as well as not requiring the donor to be moved following organ procurement (Galuta et al., 2020; Valtcheva et al., 2016). This not only reduces the postmortem interval for spinal tissue collection, reduces the number of procedures requiring institutional review board approval, but also does not interfere with funeral arrangements for the donor. We then integrated elements such as protective tissue preparation solutions and tissue sectioning parameters from our previously refined and published rodent methodology, which consistently produces high-quality slices for electrophysiology in adult rat spinal cord tissue (Dedek et al., 2019, 2022, 2024; Harding et al., 2021; Hildebrand et al., 2014, 2016), which is also technically challenging. Finally, we made several important optimizations from the above-mentioned human and rodent protocols to account for the size of the tissue, its structural integrity, and anatomical differences between species. The final procedures described here have resulted in an assay that reliably produces slices for electrophysiological examination of the hSC. These procedures include a surgical extraction technique that minimizes trauma to the spinal cord and nerve roots, a microdissection technique for preparing the hSC for sectioning, and a sectioning assay using a high-quality vibratome that ensures viability is preserved during the sectioning process. Finally, a modified recovery setup for the slices enables adequate oxygenation of the slices, with a way to track individual slices to ensure slices recover for the correct amount of time at each slice recovery stage. These studies have allowed for the comparison of both synaptic glutamatergic signalling and circuit-level activity between sex and species, thus opening the door to translational research and drug screening assays (Dedek et al., 2019, 2024).

## Applications of the method

Acute slice electrophysiology is a powerful technique that can be used to study functional neuronal outputs, including action potentials, synaptic transmission and plasticity, ion channel activity, membrane potential changes, and network activity. The foundation of our understanding of neuronal physiology is based on animal models, and until the development of this assay, cross-species comparisons directly assessing baseline neuronal physiology within spinal circuitry have not been possible (Dedek et al., 2024). Furthermore, conducting experiments using hSC tissue is an opportunity to validate pathophysiological pathways and molecular targets identified using rodent models (Dedek et al., 2019, 2022). Approaches using biophysical parameters to characterize synaptic responses (Dedek et al., 2024; Dedek & Hildebrand, 2024), can be combined with pharmacological studies for both basic science research and drug screening assays.

Beyond electrophysiology, the highly viable tissue slices prepared using this assay can be used in many other scientific applications. Tissue can be immediately flash-frozen in the operating room (OR) and used for single-nucleus RNA sequencing (Yadav et al., 2023). Additionally, mechanisms of pathological pain can be modelled *ex vivo* by incubating slices in growth factors, such as brain-derived neurotrophic factor (BDNF) (Dedek et al., 2019, 2022). Tissue can be fixed in paraformaldehyde and frozen for immunohistochemical analysis (Armstrong et al., 2022; Dedek et al., 2019, 2022; Parnell et al., 2023; Temi et al., 2021). Finally, the tissue can be flash-frozen for biochemical techniques such as western blot and quantitative PCR (Dedek et al., 2019, 2022; Murray-Lawson et al., 2024). Thus, complementary approaches can be paired with the hSC slice assay described here to ensure maximal use of the precious tissue, as well as to allow for the probing of scientific questions using multiple techniques.

## Experimental design

Overview – the tissue preparation consists of 4 distinct stages: surgical extraction of hSC tissue, tissue microdissection, sectioning and recovery, and use of slices for either patch-clamp or hdMEA recordings (Figure 1). Following the removal of organs for transplant, hSC can be extracted from consenting donors in the operating room. hSC is collected into oxygenated protective aCSF solution containing kynurenate and is immediately transferred to the laboratory. Under constant bubbling with carbogen gas, hSC tissue is then micro-dissected to remove nerve roots and meninges and is then glued onto an agarose block and specimen plate, which are then placed into the buffer tray of a vibratome. At a slow speed and wide cutting amplitude, the hSC tissue is then sectioned into 500µm slices. Slices are transferred to a custom recovery chamber in a water bath at 34°C for 40 minutes, after which slices are left to passively cool to room temperature for use in electrophysiological recordings.

**Figure 1.**
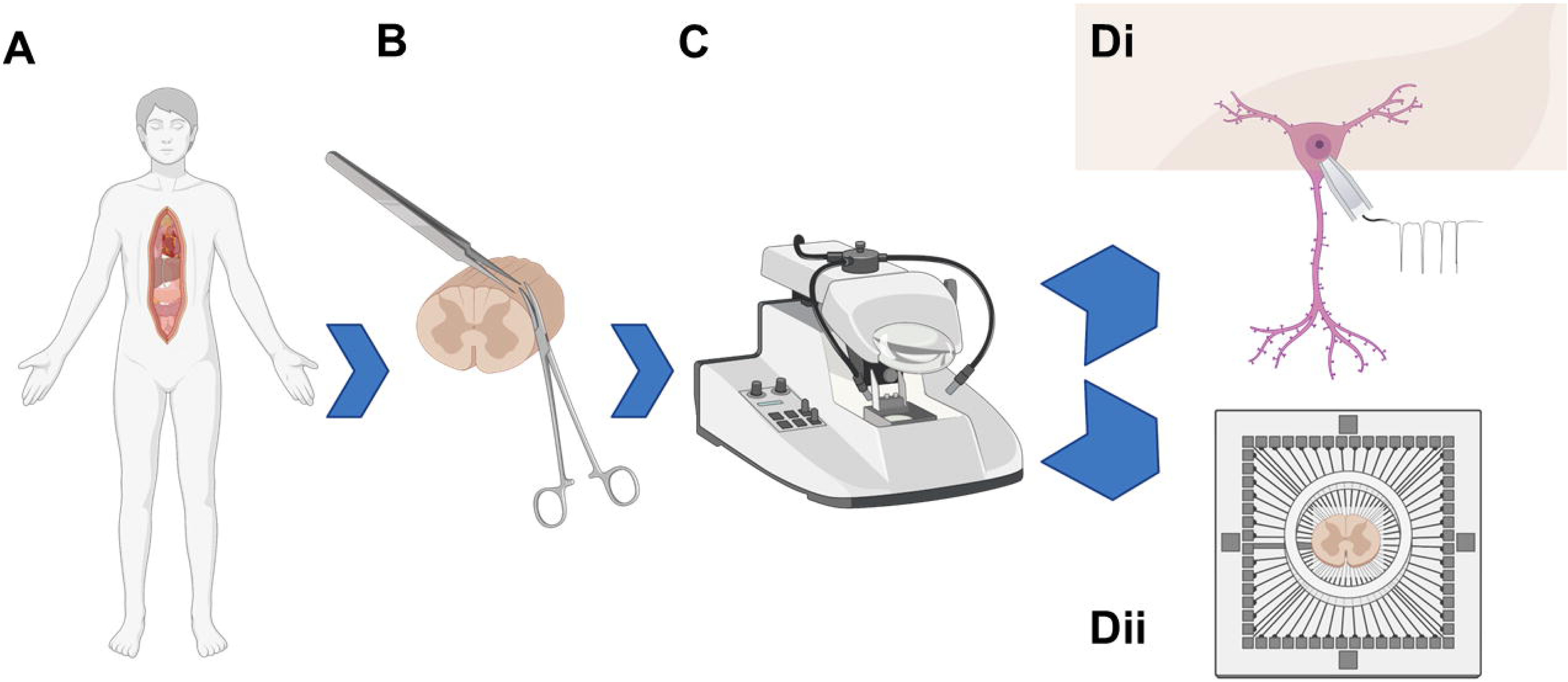
Schematic of workflow for human tissue preparation for electrophysiology recording. A) Spinal cord tissue is first harvested from organ donors. B) Tissue is then micro-dissected before C) sectioning using a vibratome. D) Spinal cord slices are then ready for electrophysiological analysis using either Di) patch-clamp or Dii) multielectrode array recording. Figure created with BioRender.com

Expertise needed to implement the protocol – At many institutions/hospitals, a licensed surgeon is required to oversee the tissue collection. Once tissue is collected, a graduate student or postdoc with experience in acute slice electrophysiology has the required skills to perform this protocol. To maximise the use of the acute tissue, it is suggested multiple team members share the tissue for parallel, concurrent electrophysiological experiments.

Access to human organ donor tissue – collection of hSC tissue is contingent on a collaboration with a transplant service or organ procurement organisation (depending on the structure of the medical system where tissue is collected). After approaching the transplant service to establish a collaboration, appropriate regulatory approval and documentation are required, such as IRB approval, consent forms for family, and establishing a protocol for the documentation required for consent. The specific protocol for access to the operating room, surgical tools available, and the logistics for timing and notification will vary between institutions and must be established with the transplant service and institution. Since establishing our human tissue pipeline in 2017, we have received tissue on average 10 times/year, barring a temporary pause in tissue collection during the outbreak of COVID-19, at a trauma center serving a population of approximately 2 million people. At our institution, we are notified once a donor’s family consents to collection for our study. An operating room for organ collection is then scheduled, typically 24-48 hours in advance.

## Limitations

As with any acute slice preparation, several factors may pose problems in the preparation of high-quality slices. Factors out of experimental control, such as the overall condition of the donor before undergoing organ harvest, as well as possible delays in extracting organs for transplant, may negatively affect the tissue. In some cases where few organs are collected, surgical extraction of the spinal cord may be difficult, due to lack of space in the body cavity to access the spinal column via an anterior approach. In such cases, it is likely only 3-5 centimetres of hSC can successfully be harvested, when up to 30 centimetres can be extracted when all possible organs are harvested. We have found that tissue from all ages of donors can yield high-quality tissue, but donors who spent less time (<5 days) in intensive care before undergoing organ collection are more likely to produce high-quality tissue than those who were in ICU for an extended time. Furthermore, successful electrophysiological experiments can be performed from neurologic determination of death (NDD) and determination of cardiac death (DCD) donors; however, the shortened length of time between removal of supportive care and tissue collection in NDD cases yields more reliably high-quality tissue and a higher success rate for electrophysiological experiments. As is the case with all acute slice preparations, axotomy induces an inflammatory response and some cellular loss during the slice preparation is inevitable. The slice recovery process outlined here, paired with viability validation using TTC staining ensures tissue that is used for experiments is robust and of sufficient quality.

## MATERIALS

### Reagents

#### Tissue from human organ donors

- Extraction procedures and consent forms were approved by the Internal Review Board (IRB)
- Tissue should be extracted as soon as possible to minimize the postmortem interval; Typically extract spinal cord tissue 1h-3h postmortem, immediately following surgical resection of donor organs for transplantation
- We have detected spikes/spontaneous activity with tissue extracted up to 4h postmortem
- Take note of time of life-support withdrawal and cross-clamp time

- > 96% Ethanol
- Carbogen gas (95% O^2^, 5% CO^2^)
- Deionized water
- TTC (Sigma-Aldrich, cat. no. T8877-10G)

- Warning - flammable; can cause eye/skin irritation
- Agarose (Sigma-Aldrich, cat. no. A6013-100G)
- Instant adhesive (Loctite, part no. 46551)

- Warning - Combustible liquid, causes eye irritation, may cause respiratory irritation and genetic defects
- Protective aCSF

- Sucrose (Sigma-Aldrich, cat. no. S9378-1KG)

▪ Warning - may form combustible dust concentrations in air
- NaCl (Sigma-Aldrich, cat. no. 71380-1KG)
- D-(+)-Glucose (Sigma-Aldrich, cat. no. G8270-1KG)
- NaHCO_3_ (Sigma-Aldrich, cat. no. S5761-500G)
- KCl (Fisher, cat. no. P217-500)

▪ Warning - causes eye irritation; may cause respiratory tract irritation
- NaH_2_PO_4_ (Sigma-Aldrich, cat. no. S0751-500G)
- CaCl_2_ dihydrate (Fisher, cat. no. BP510-100)

▪ Warning - causes serious eye irritation
- MgSO_4_ anhydrous (Fisher, cat. no. M65-500)
- Kynurenic acid (Sigma-Aldrich, cat. no. K3375-5G)
- Slice External Recording Solution

- NaCl (Sigma-Aldrich, cat. no. 71380-1KG)
- KCl (Fisher, cat. no. P217-500)

▪ Warning - causes eye irritation; may cause respiratory tract irritation
- NaHCO_3_ (Sigma-Aldrich, cat. no. S5761-500G)
- NaH_2_PO_4_ (Sigma-Aldrich, cat. no. S0751-500G)
- CaCl_2_ dihydrate (Fisher, cat. no. BP510-100)

▪ Warning - causes serious eye irritation
- MgCl_2_ hexahydrate (Fisher, cat. no. BP214-500)

▪ Warning - irritating to eyes and respiratory system
- D-(+)-Glucose (Sigma-Aldrich, cat. no. G8270-1KG)
- Internal Recording Solution

- Gluconic acid (Fisher, CAS no. 526-95-4)

▪ Warning - causes severe skin burns and eye damage
- CsOH monohydrate (Fisher, CAS no. 35103-79-8)

▪ Warning - causes severe skin burns and eye damage.
- CsCl (Fisher, CAS no. 7647-17-8)

▪ Warning - reproductive toxicity, suspected of damaging fertility or the unborn child
- BAPTA, 1,2-bis 2-aminophenoxy ethane-n,n,n’,n’-tetraacetic acid, (Thermo Fisher Scientific, CAS no. 85233-19-8)
- HEPES (Fisher, cat no. BP310-500)

▪ Warning - may cause respiratory irritation
- Mg-ATP (Sigma-Aldrich, cat. no. A9187-1G)

▪ Warning - may cause damage to organs
- Na_2_-GTP (Sigma-Aldrich, cat. No. G8877-100MG)

### Equipment

#### Surgical Spinal Cord Extraction

- Orthopedic mallet (Blacksmith Surgical, cat. no. BS-13-34011)
- Stryker System 7 Sternal Saw (not autoclavable, cat. no. 7207-000-000)
- 32-mm straight tip bone osteome (Blacksmith Surgical, BS-13-34329)
- Debakey tissue forceps (Sklar Surgical Instruments, cat no. 52-5307)
- Straight mayo scissors (Sklar Surgical Instruments, cat no. 15-1555)
- Metzenbaum scissors (Sklar Surgical Instruments, cat no. 22-1507)
- Harrington-mixter clamp (Sklar Surgical Instruments, cat no. 55-3012)
- Scalpel handle #3 (BS-01-10001) with no.10 blades (Bard-Parker, cat. no. 371110)
- 50-ml Conical tubes (Greiner, item no. 210270)
- Container filled with ice (sufficient to accommodate 6 conical tubes) Spinal Cord Microdissection
- Stereo microscope (Leica, M50)
- Dumont #5 straight tip forceps (Fine Science Tools, item. no. 11251-10)
- Scalpel handle with no. 10 blades (Bard-Parker, cat. no. 371110)
- Debakey tissue forceps (Sklar Surgical Instruments, cat no. 52-5307)
- Straight edge Vannas microdissection scissors (World Precision Instruments, cat. no. 500086)
- 50 mL Petri dishes (VWR)
- Styrofoam tray

#### Tissue Sectioning

- Vibratome with Vibrocheck (Leica, VT1200S)
- Water Bath (Fisher Scientific, Isotemp 2340)
- Custom glass dropper
- 1L Beaker
- 300mL petri dish (VWR)
- 2x Custom slice holder, or commercially available large slice holder

#### Patch-Clamp Recordings

- Flaming/brown micropipette puller (Sutter instrument, P-97)
- Microforge (Narishige, MF-83)
- Borosilicate glass capillary tubes (1.5/1.17mm outer/inner diameter) (Sutter instrument, item no. BF150-117-10)**
- Microscope Axio Examiner.A1 (Zeiss)
- Headstage amplifier (Molecular Devices, CV-7B)
- Amplifier (Molecular Devices, Multiclamp 700B)
- Digitizer (Molecular Devices, Digidata 1550)
- Infrared Microscopy Camera (Dage-MTI, IR-1000)
- Micromanipulator (Scientifica, PS-7500)
- Peristaltic pump (Fisher, CTP300)
- Tygon tubing, assorted sizing (Fisher, cat no. 14-179-110)
- Tissue anchor (Warner Instruments)
- Optical Table (ThorLabs)

#### High-Density Microelectrode Array (hdMEA) Recordings

- hdMEA system (3Brain, BioCAM X)
- Peristaltic pump (Fisher, CTP300)
- Tissue anchor (3Brain)
- Tygon tubing, assorted sizing (Fisher, cat no. 14-179-110)
- Blunt-fill 16G needles (Fisher, cat no. BD 305180)

#### Software

- BrainWave v.5 (3Brain)
- Clampfit v.11.2 (Molecular Devices)

#### *See equipment set-up

#### **!Critical step

##### Reagent Set-Up 5% Agarose

5% (w/v) solution in protective aCSF, heated to dissolve agarose and refrigerated to set in a petri dish. The depth of the agarose gel should be 1 centimetre.

##### Protective aCSF

To prepare 1L of protective aCSF, combine the reagents listed in the table below in deionized water. Store at 4°C and use within 2 days. Immediately prior to use, allow to bubble with carbogen for a minimum of 20 minutes.

| Reagent | Final Concentration (mM) | Molecular Weight (g/mol) | 1L |
| --- | --- | --- | --- |
| Sucrose | 50 | 342.3 | 17.115g |
| NaCl | 92 | 58.44 | 5.376g |
| glucose | 15 | 180.156 | 2.702g |
| NaHCO<sub>3</sub> | 26 | 84.007 | 2.184g |
| KCl | 5 | 74.5513 | 0.373g |
| NaH<sub>2</sub>PO<sub>4</sub> | 1.25 | 119.98 | 0.172g |
| CaCl<sub>2</sub> | 0.5 | 110.98 | 0.074g |
| MgSO<sub>4</sub> | 7 | 120.366 | 0.843g |
| Kynurenic acid* | 1* | 189.17 | 0.039g* |

\*Dissolve 0.039g of kynurenic acid into 200mL of protective aCSF the day of experiments
NOTE: freeze extra protective aCSF with kynurenate in an ice cube tray in advance of hSC experiments. To help keep the preparation cold during tissue extraction, microdissection, and sectioning, ensure some protective aCSF ice is always visible at all stages up until slice recovery.

##### Slice External Recording Solution (aCSF)

To prepare 1L of slice external recording solution, combine the reagents listed in the table below in deionized water. Store at 4°C and use within 1 week. Immediately prior to use, allow to bubble with carbogen for a minimum of 20 minutes.

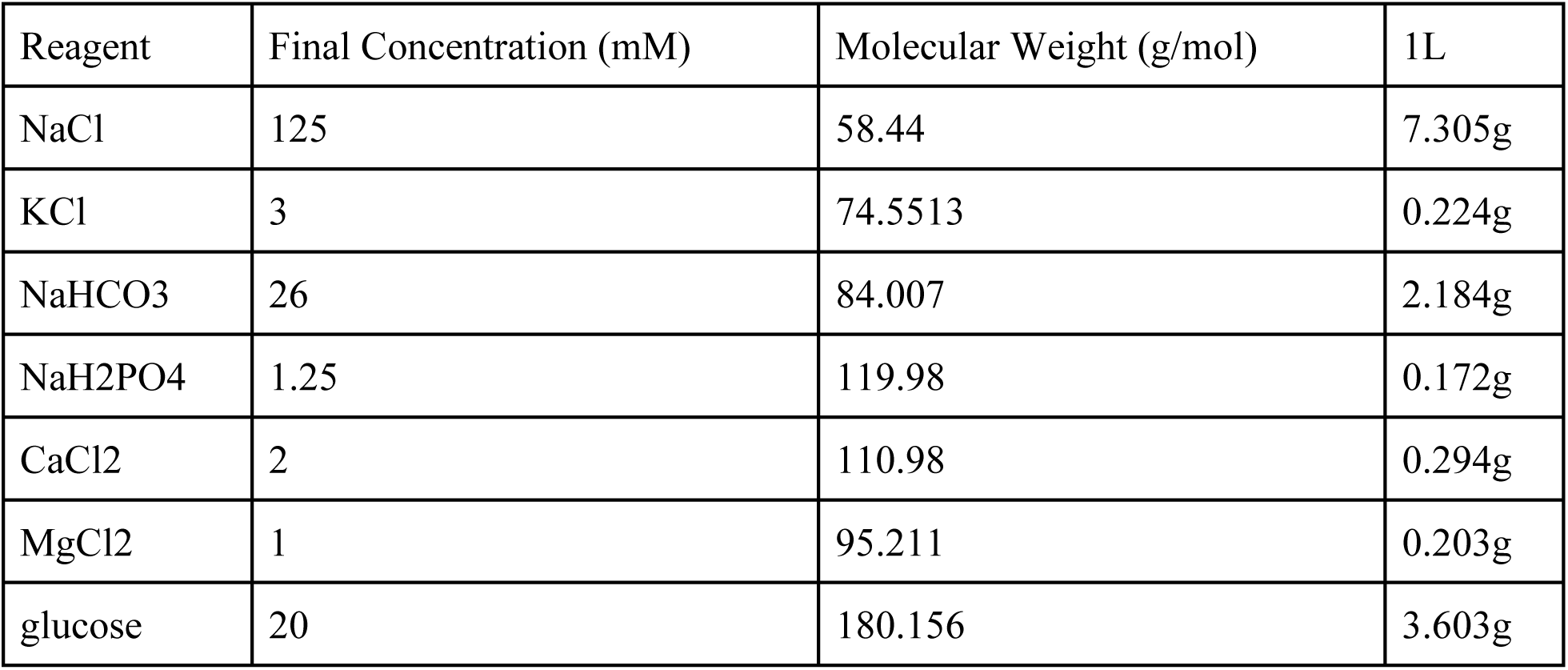

##### Internal Recording Solution

To prepare 10mL of internal recording solution, combine all reagents except for Mg-ATP and Na2-GTP in 9.5mL of deionized water. Adjust the pH to 7.25 with CsOH and check the volume to ensure it amounts to 10mL. Add Mg-ATP and Na2-GTP. 295 mOsm. Filter then freeze 1mL aliquots and store at -20°C and use within 6 months.

| Reagent | Final Concentration (mM) | Molecular Weight (g/mol) | g/10mL |
| --- | --- | --- | --- |
| Gluconic acid | 105 | 196.16 | 0.206 |
| CsOH | 105 | 149.912 | 0.157 |
| CsCl | 17.5 | 168.36 | 0.029 |
| BAPTA | 10 | 380.35 | 0.038 |
| HEPES | 10 | 238.3012 | 0.024 |
| Mg-ATP | 2 | 529.47 | 0.0101 |
| Na<sub>2</sub>-GTP | 0.5 | 523.18 | 0.0026 |

##### TTC Staining Solution

Prepare 10 mL of a 2% w/v solution of TTC in room temperature PROTECTIVE ACSF without kynurenate that has been bubbled with carbogen.

#### Equipment Set-Up

##### Sternal Saw

Ensure the blade of the sternal saw is facing away from the handle, so that the saw cuts when pushing the saw away from you. Often this required turning the blade 180°.

##### Vibratome Calibration

Securely attach the vibrating microtome blade and follow Vibrocheck calibration instructions as described in the manufacturer’s protocol. Ensure a new blade is installed and the system is calibrated. This calibration is critical to ensure that the vertical deflection of the blade is minimized, thus producing slices with viable cells on the surface of the slice. **

##### Custom Glass Dropper

Modify a standard glass Pasteur pipette by scoring the taper of the pipette with a diamond-tip glass cutter. Then break the tip off of the pipette and attach the bulb to the cut end. A glass dropper gives more precision when transferring slices between solutions and reservoirs, and tissue is less likely to attach to it.

##### Custom Slice Holder

Cut 1-inch polyvinyl chloride (PVC) piping and using hot glue, assemble them together to form a 9-unit grid. Secure the plastic screen beneath the grid using the same adhesive (See Supplementary Figure 1). Commercial options are also available.

##### Water Bath for Slice Recovery

Place a 1L beaker filled with approximately 400mL PROTECTIVE ACSF (without kynurenate) in a water bath. Place the custom slice holder in the beaker; it will float just below the surface, ensuring slices are separated and immersed in PROTECTIVE ACSF, without sitting on the bottom of the beaker and risking having an area that is not exposed to fresh oxygenated solution. Heat the water bath so that the temperature of the PROTECTIVE ACSF is exactly 34℃. Bubble the PROTECTIVE ACSF continuously with carbogen.

##### Patch-clamp Rig and hdMEA

Ensure all components of the equipment are on, aCSF solution is being bubbled with carbogen in the perfusion reserve, and recording software is open.

##### hdMEA Chip Preparation

If required based on manufacturer recommendations, 3 days prior to recordings fill the chip chamber with PBS to hydrate the chip. Immediately prior to use, clean the contact pads of the chip with 96% ethanol and allow time to dry. Rinse the reservoir once with ethanol and twice with distilled water.

##### Recording Glass Pipette Preparation

To pull the patch pipette, mount the borosilicate glass capillary tube onto the puller as described in the manufacturer’s protocol. Using the microforge, gently fire-polish the glass patch-clamp pipettes, with 6-12 MΩ resistance once filled with internal. Ensure the pipette has a tapered and conical structure to facilitate sealing and prevent plugging.

###### Spinal Cord Extraction (Timing - 30-50 minutes)

**CRITICAL** The amount of space within the visceral cavity to perform the following steps will vary depending on the organs harvested for donation, as well as the amount of visceral body fat of the donor. The procedure is made more difficult with fewer organs removed and/or if the donor has a large amount of visceral fat. In the case of limited access to the spinal column, extend as rostrally as possible. The dissection tools used below were made available for this extraction protocol in the operating room. Investigators should coordinate with their transplant service and hospital regarding whether they need to provide their own instruments.

**! CAUTION** Sterile operating room procedures must be followed for tissue extraction in the operating room. Exercise proper safety precautions (e.g. gloves, mask, eye protection, scrubs, surgical gown) when working with human tissues.

1. Immediately before leaving the laboratory space to go to the operating room, prepare several 50mL conical tubes filled with bubbled 0-4℃ protective aCSF containing kynurenate in a Styrofoam box filled with ice. Ensure that the canonical tubes have screw-top lids, which are securely tightened to avoid dissipation of oxygen. The remaining protective aCSF should continue to be bubbled on ice in the laboratory space during the spinal cord extraction.
2. Using a surgical towel and retractor, contain the remaining organs to expose the spinal column.
3. Locate the sacral promontory. Then, count the lumbar vertebrae to identify L2.
4. Use an osteotome and mallet to make a transverse, wedge-shaped osteotomy through the L2 vertebral body (Figure 2Ai). Ensure the wedge is sufficiently wide to expose the spinal canal and allow the footplate of the sternal saw to be placed inside the spinal canal, without penetrating the dura.
5. Mobilize all organs to one side, exposing as much length of the spinal column as possible on one lateral side. Insert the footplate of the sternal saw and angle the saw at 45° medially (Figure 2B). Pushing the sternal saw away from you, cut through the vertebral bodies in a caudal to rostral direction as rostrally as the accessibility within the body cavity will allow (Figure 2Aii). In cases where the heart and lungs are removed, this may be to the top of the surgical incision in the ribcage. If many organs remain, cut as high as possible. Then move the organs to the other side and repeat on the other side.

**Figure 2.**
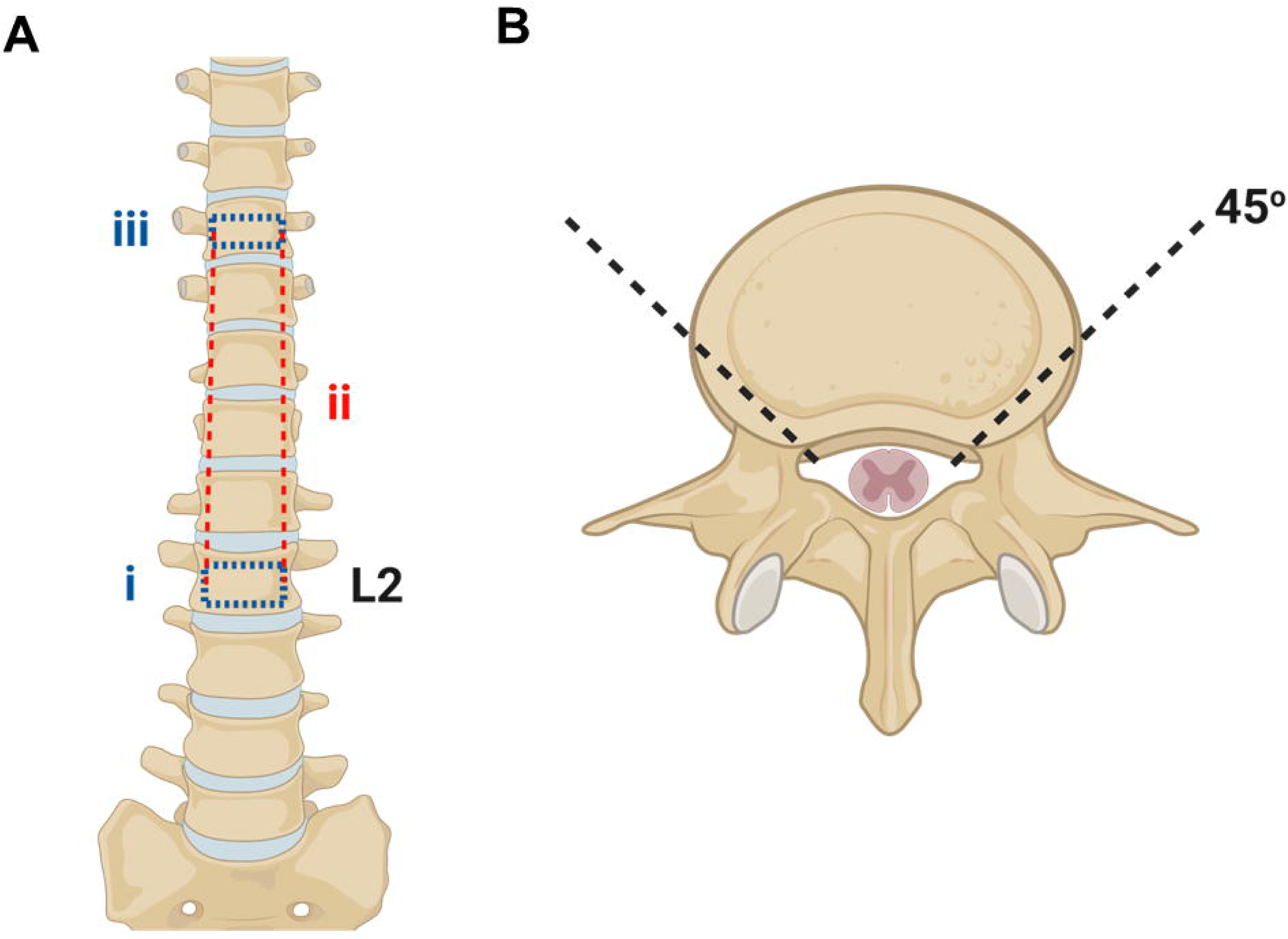
**Surgical approach for spinal cord extraction**. A) i: Using a straight, wide osteotome and surgical mallet, the L2 vertebral body is transected, stopping at the spinal canal. ii: the two incised vertebral bodies are connected by an incision made using a sternal saw. iii: The most superior vertebral body that is accessible is transected to allow for the removal of the section of vertebral disk and expose the spinal canal. B) A 45-degree angle must be used in ii, ensuring the sternal saw clears the vertebral bodies and without damaging the spinal cord beneath. Figure created with BioRender.com

CRITICAL STEP Once inserted into the spinal canal, it is critical to hold the footpad of the saw firmly against the anterior wall of the spinal canal to ensure the footpad does not damage the spinal cord below.

CRITICAL STEP Do not change the angle of the sternal saw. Changing this angle may break the blade of the saw or result in difficulty retrieving the saw.

##### ? Troubleshooting (Table 1)

6. Use a straight osteotome to detach the vertebral bodies, taking care not to perforate the dura or damage the spinal cord below (Figure 2Aiii).
7. Using a closed Harrington-Mixter clamp to gently lift the thecal sac, transect the thecal sac with Mayo or Metzenbaum scissors at the L2 level. Holding only the dura (not any roots or the hSC), gently lift the thecal sac. Using Mayo or Metzenbaum scissors, transect the nerve roots and fascia that hold the thecal sac in the spinal canal. Once the exposed length of the thecal sac is freed, transect the rostral end of the exposed thecal sac. Transfer to a folded surgical towel for microdissection.

CRITICAL STEP Do not lift the dura more than 20° from the spinal canal, and do not put excess tension on the thecal sac. Lifting the thecal sac at too abrupt an angle will severely damage the grey matter of the hSC. It is better to carefully cut under and around the thecal sac without fully being able to see than to lift it too high.

8. On a surgical towel, use forceps to lift the dura off the spinal cord. Using Mayo scissors, quickly and gently cut from caudal to rostral, exposing the hSC.
9. Using forceps and a number 10 blade, section the hSC into 1.5cm pieces and immediately place in prepared, ice-cold, pre-bubbled protective aCSF with kynurenate for transport to the laboratory.

**Table 1.** Troubleshooting Guide

| <u>Step</u> | <u>Problem</u> | <u>Possible Reason</u> | <u>Solution</u> |
| --- | --- | --- | --- |
| 5. | Sternal saw cannot span the vertebral bodies | Large vertebral body diameter | Use a straight osteotome to perform the procedure. This adjustment may be necessary, but significantly increases the time required for hSC extraction. |
| 18. | Slices are tearing or lack integrity | This could happen for a number of reasons – please match possible reason with correspondingly numbered solution.<br>1. Tissue is of insufficient quality to slice<br>2. Meninges are catching the blade and tearing tissue<br>3. Glue has wicked in front of the blade and is obstructing the blade | 1. The tissue may be damaged due to the circumstances surrounding death of the donor, or because of issues with the tissue extraction. Review extraction to ensure that no force was put on the hSC sample during extraction.<br>2. If another tissue sample is available, restart the procedure with a new hSC segment and ensure to carefully remove ALL roots and meninges.<br>3. If another hSC tissue sample is available, restart and ensure a minimal amount of glue is<br>used to secure the hSC on the vibratome chuck. |
| 20. | Slice is not changing colour to red following TTC staining | The sample is damaged, and no viable cells remain. | Carefully examine all previous stages to determine if something could have damaged the hSC sample. If no reason is identified, it may be due to the circumstances surrounding the death of the donor. |
| 23. | Cracks emerge in the hSC sample while it is mounted to the specimen plate | Damage during extraction or microdissection, incorrect sectioning parameters, required more agarose support. | If tears or cracks appear in the hSC tissue sample while it is being sliced, increase the thickness of the slice to get past the damaged area. Slow down the cutting speed and ensure a cutting blade amplitude of 2.75mm is being used. Ensure the tissue sample is being supported by the agarose block. |
| 25-31 | No activity in recordings | Slices may no longer be viable | Repeat Step 20 (TTC staining) to check for slice viability with one of the remaining slices. If no colour change is observed, end experiments, as the tissue is no longer viable. |

CRITICAL STEP Time sensitive. It is critical to get the extracted tissue into oxygenated protective solution as quickly as possible; It is critical that the number 10 blade is used gently. Unlike typical scalpel use, we recommend using the forceps to gently secure the hSC on either side, not putting any pressure on the hSC itself. Then, using only the weight of the number 10 blade and handle, slowly saw back and forth across the hSC. Using the typical scalpel technique of one swift incision creates excess shearing force and downward pressure and can be very damaging to the hSC.

TIP: We have found that using the conus terminalis/sacral spinal cord region for electrophysiological recordings is the most practical, as the smaller tissue diameter makes sectioning easier.

##### Spinal Cord Microdissection (Timing – 30-60 minutes)

**! CAUTION** Follow proper safety precautions when working with and disposing of materials containing human tissues.

10. Place a piece of hSC in a 100mm petri dish filled with ice-cold protective aCSF with kynurenate to fully submerge the hSC. Keep the petri dish on ice in a Styrofoam tray for the duration of cleaning and dissection. Bubble the protective aCSF with carbogen and continually add protective cutting solution ice cubes as they melt over time.
11. Using Dumont #5 straight-tip forceps and straight-edge Vannas microdissection scissors, gently remove the anterior and posterior roots. Begin by securing a root with the forceps, and with the scissors gently follow the root to the surface of the hSC. Gently, using as little force as possible, align the scissors so they are parallel to the surface of the hSC and cut to remove the root (Figure 3A, Supplementary Video 1). Repeat for all roots.
12. Using Dumont #5 straight-tip forceps and straight-edge Vannas microdissection scissors, pinch the remaining meninges as superficially as possible and cut to remove a small portion of the meninges. Continue this process piece by piece until all the meninges are removed (Supplementary Video 1).

**Figure 3.**
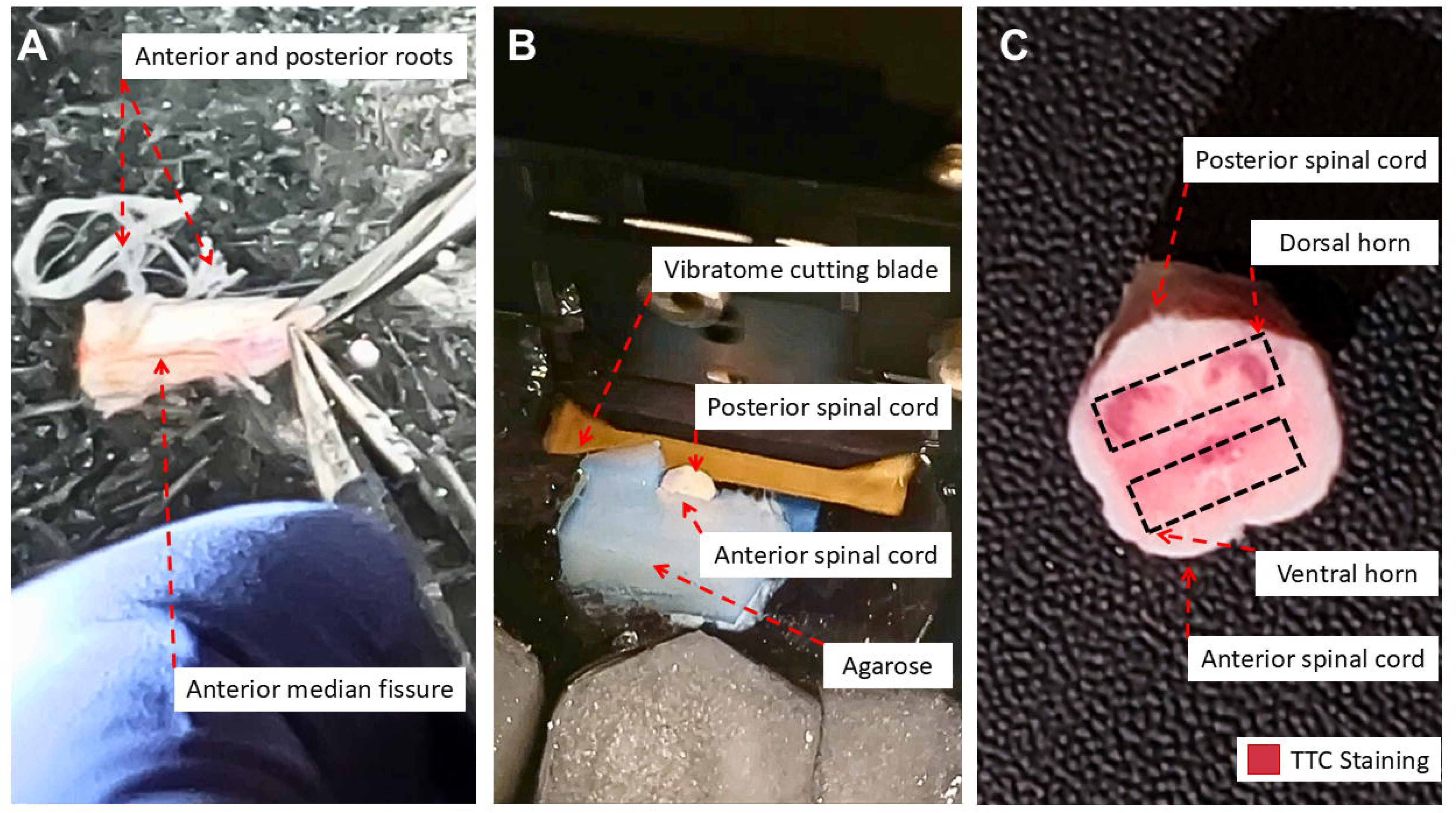
Spinal cord preparation. A) Tissue microdissection: it is critical to remove all roots, followed by all meninges prior to sectioning. B) Sectioning tissue on a vibratome, using blocks of agarose to support the spinal cord. The spinal cord is secured using glue on the anterior side for examination of the posterior horn (equivalent to the dorsal horn in rodents). C) the first slice removed during sectioning is placed in a 2% 2,3,5-Triphenyltetrazolium chloride (TTC) solution for 30 minutes as a readout of tissue viability before proceeding with experiments.

CRITICAL STEP This step is painstaking and slow, but the utmost care should be taken to remove as much of the meninges as possible to ensure sectioning on the vibratome does not damage the slices. We have found that the more difficult the removal of the remaining meninges, the higher quality of hSC. If the pia and arachnoid layer easily peel away, or if there is excessive fraying of the underlying white matter, this may indicate the hSC is disintegrating and not viable for experiments. To fully remove the remaining meninges, it is often necessary to pinch and cut away a small amount of the superficial white matter. This does not impact the underlying grey matter, which is located more internally than in the rodent spinal cord.

13. Perform a visual inspection of the exposed area of the grey matter to ensure that there are no hematomas or other damage and that the edge of the tissue is straight and perpendicular to the length of the hSC. If the edge is not straight or there is damage, use Debakey tissue forceps and a number 10 blade to remove tissue and straighten out the edge of the piece of hSC, as described in Step 9.

##### Spinal Cord Sectioning and Recovery (Timing – 2-4 hours)

**! CAUTION** Follow proper safety precautions when working with and disposing of materials containing human tissues.

14. Using a razor blade, cut a piece of agarose that is barely longer and wider than your hSC segment (typically 12mm x 14mm). Ensure the agarose has approximately 90° corners to make sure the slices will be straight. Glue one of the two smallest faces of the agarose block to a vibratome specimen plate using instant adhesive (Figure 3B, Supplementary Video 1). Confirm that the angle created by the chuck and the agarose block is 90°.
15. Place a small dab of instant adhesive on the specimen holder directly in front of the agarose block. Place a thin line of adhesive on the edge of the agarose that faces the first dab of adhesive. With a folded kimwipe, spread the adhesive into a thin, even coat.

CRITICAL STEP ! Do not use excessive adhesive. Excess adhesive may wick around the hSC when solution is poured in and then may catch the vibratome blade, damaging the slices.

16. Using Debakey forceps and the flat end of a scoopula, pick up the hSC segment by the posterior side if studying the posterior (dorsal in rodents) horn, and by the anterior side of studying the anterior (ventral in rodents) horn. Gently touch the edge of the hSC segment to the edge of the petri dish to allow excess protective aCSF to flow off of the hSC segment. Gently place the hSC segment on the specimen holder and agarose block covered in adhesive, with the area of interest (anterior or posterior horns) facing out (Figure 3B, Supplementary Video 1).
17. Immediately place the specimen holder with the attached hSC segment in the buffer tray of a vibratome, with the ice tray lined with ice to ensure it remains ice-cold throughout sectioning. Immediately immerse in ice-cold protective aCSF with kynurenate bubbled with carbogen, by pouring the solution gently into the side of the buffer tray until it reaches the height to completely submerse the spinal cord segment. Bubble with carbogen throughout sectioning and continually add protective cutting solution ice cubes to maintain a temperature of 0-4°C.
18. At a cutting speed of 0.01-0.02mm/s and a horizontal blade amplitude of 2.75mm, remove a thick (>500 μm) slice off the mounted hSC. This slice should be thick enough to create a level surface from which future 500μm slices can be cut. A number 10 scalpel blade may be required to free the slice from the agarose block (Supplementary Video 1).

##### ? Troubleshooting (Table 1)

19. While this first slice is being sectioned, prepare the TTC solution.
20. Place the freshly sectioned thick hSC slice in a well of a 6-well plate, immersed in TTC solution. Place in a cell-culture incubator for 30 minutes. Continue with sectioning while the slice is incubating in TTC. After 30 minutes, check to see if a red colour change has occurred throughout the grey matter, indicating mitochondrial activity and slice viability (Hatfield et al., 1991) (Figure 3C). If the slice does not change colour to red, the tissue is not viable for electrophysiological recordings. Try once more with another slice, and if no colour change occurs once more, end the process here.

##### ? Troubleshooting (Table 1)

21. After the first thick hSC slice, section the hSC at 500µm. When the first 500µm slice is ready, release it from the agarose block using a number 10 blade and use the custom dropper to transfer the slice to the prepared custom slice recovery chamber (Supplementary Figure1) in the heated water bath.
22. Set a running timer. Continue sectioning and placing slices sequentially in the slice recovery chamber. 40 minutes after the first slice was put in the slice recovery chamber, use a motorized pipettor to remove approximately 150mL of 34℃ protective aCSF from the 1L beaker and place it in a 300mL petri dish containing a second, smaller custom slice holder. Move the first slice from the first chamber in the second slice holder, and ensure this solution is bubbled with carbogen. This 300mL petri dish will passively cool to room temperature in approximately 30 minutes, after which the slice can be used for experiments.
23. Continue sectioning. After transferring the first slice out of the water bath slice recovery chamber, move one additional, sequential, slice from the water bath slice recovery chamber to the room temperature recovery using the custom dropper after each new slice is finished on the vibratome. Note the approximate time it takes for one slice to finish sectioning.

##### ? Troubleshooting (Table 1)

24. Once you are finished sectioning on the vibratome, set a timer for the amount of time it took to complete sectioning a slice. When this timer is up, move the next-oldest slice from the water bath recovery to the room temperature recovery. Ensure slices remain at room temperature recovery for 30 minutes before starting experiments.

Note: If you have multiple team members working, you can begin electrophysiology experiments while one team member continues sectioning and managing slice recovery.

##### Electrophysiological Recording

**! CAUTION** Follow proper safety precautions when working with and disposing of materials containing human tissues.

**Note:** If enough team members are present and you have both a patch-clamp rig and hdMEA, Option A and Option B can run concurrently.

**? TROUBLESHOOTING** If recordings are not yielding activity, or cease to yield activity, repeat Step 20 (TTC staining) to check for slice viability with one of the remaining slices. See Table 1.

##### Option A: Patch-clamp Recording (Timing – 2-8 hours)

25. Pick up a single slice using the custom dropper. Place it in a small petri dish containing some room temperature, bubbled protective aCSF without kynurenate. Using straight-edge Vannas dissection scissors, cut the slice in half to separate the left/right halves of the hSC. Note, if the slice is small enough in diameter to fit in the microscope well (Figure 4A) beneath your tissue anchor, this step can be skipped.
26. Place the hSC hemisection in the immersion chamber, filled with bubbled aCSF, of the patch rig. Secure with tissue anchor, and continuously perfuse with fresh, bubbled solution throughout experiments.
27. Identify your region of interest under brightfield optics. The substantia gelatinosa can serve as a useful marker. Note that the anatomy of the hSC differs substantially from rodent spinal cord (Toossi et al., 2021). Refer to an atlas if needed (Sengul et al., 2013).

##### Option B: hdMEA Recording (Timing – 2-8 hours)

28. Depending on the size of the active recording area on your hdMEA, your hSC slice will need to be cut to fit into this area. Our preparation uses a 6mm x 6mm recording area, thus hSC cut into ¼ sections fit well onto the chip. Follow Step 25 above to cut the tissue containing your area of interest to the correct size for your chip.
29. Couple the tissue to the hdMEA chip. Using a P200 pipettor, remove all solution from the chip. Place tissue anchor to secure the slice and immediately pipette on 30µL aCSF.
30. Remove the aCSF and then immediately replace on the slice, repeating this step three times, thus coupling the slice to the active recording area of the chip.
31. Continuously perfuse at a rate of 1-4 mL/minute with fresh, bubbled solution throughout experiments. Before beginning an experiment, allow the slice to acclimate on the chip for 15 minutes.

#### Timing

**Reagent Setup:** agarose preparation, internal recording solution preparation, slice external recording solution (aCSF): 3.5 hours (can be made in advance and kept on hand)

**Figure 4.**
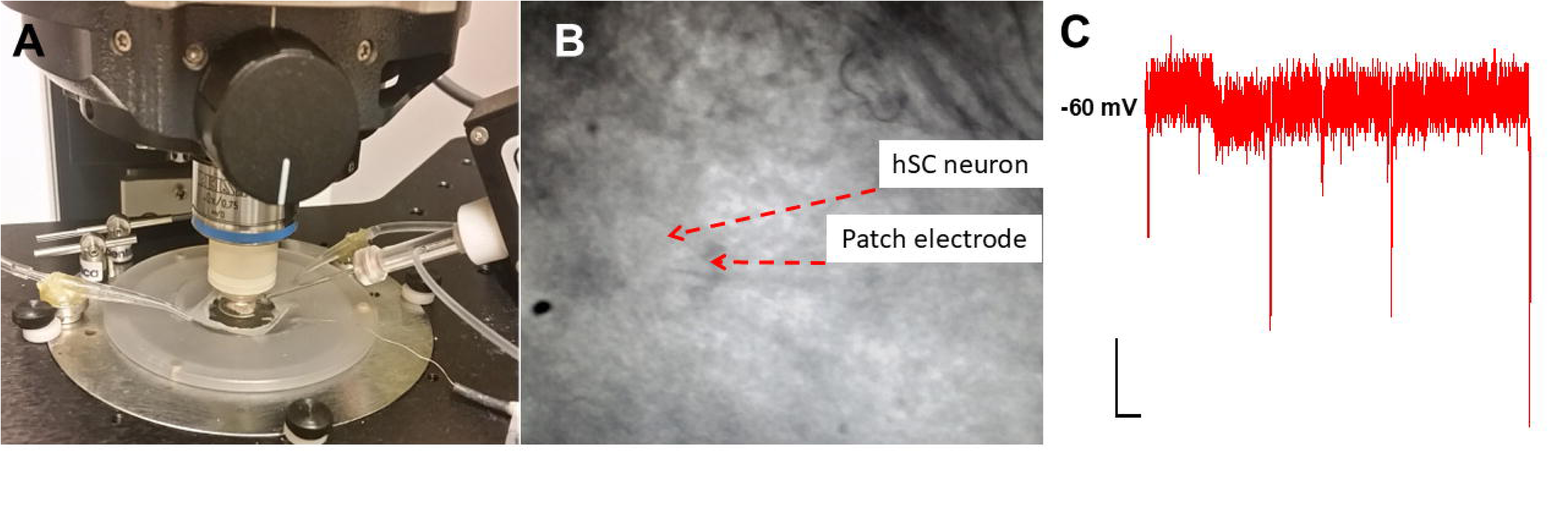
Patch clamp recording in human spinal cord slices. A) A human spinal cord slice is secured with a tissue anchor for recording. B) A patch electrode sealed onto a human superficial posterior horn neuron. C) Representative trace of postsynaptic activity at -60 mV. Scale bars = 1000 ms (x axes), 10 pA (y axes)

**Reagent setup:** make protective aCSF, cool, dissolve kynurenate, bubble protective aCSF: 2 hours

**Equipment Setup:** 1.5 hours

**Steps 1 - 9:** hSC extraction: 30 – 50 minutes

**Steps 10 – 13:** Spinal cord microdissection: 30 – 60 minutes

**Steps 14 – 24:** Tissue sectioning and recovery: 2 – 4 hours

**Steps 25 – 31:** Electrophysiological recording: 2 – 8 hours

**Troubleshooting:**

**Anticipated results:**

### Option A: Patch-clamp Electrophysiology

This protocol can be used to prepare tissue for patch-clamp recording (Dedek et al., 2019, 2024). We patched pain-processing neurons from lamina I of the superficial dorsal horn (SDH; anatomically the superficial posterior horn in humans), which was identified as being within 50 μm medially of tracts that run along the outer edge of the substantia gelatinosa. All slices used had a clear and bright substantia gelatinosa under brightfield optics. Neurons that were selected to be patched had smooth surfaces free of blebs and had lightly defined edges (Figure 4B). The criteria for recording neurons included an access resistance under 30 MΩ and leakage currents no greater than −100 pA at a holding potential (V_h_) of −60 mV (Hildebrand et al., 2014).

Whole-cell patch was established at −60 mV, allowing for the recording α-amino-3-hydroxy-5-methyl-4-isoxazolepropionic acid (AMPA) receptor-mediated miniature excitatory postsynaptic currents (mEPSCs) (Figure 4C). AMPAR mEPSC responses from 6 neurons recorded across 4 male and 2 female donors show consistent waveform characteristics (Figure 5Ai). Measuring the average amplitude (18.44 ± 8.93 pA), 10–90% rise time (0.27 ± 0.26 ms), and decay constants (2.76 ± 1.78 ms) across the sampled neurons provided insights into the biophysical properties of these excitatory synaptic responses in the human SDH at −60 mV (Figure 5Aii-iv), with faster activation and deactivation kinetics compared to that previously observed for rat lamina I neurons (Hildebrand et al., 2014).

**Figure 5.**
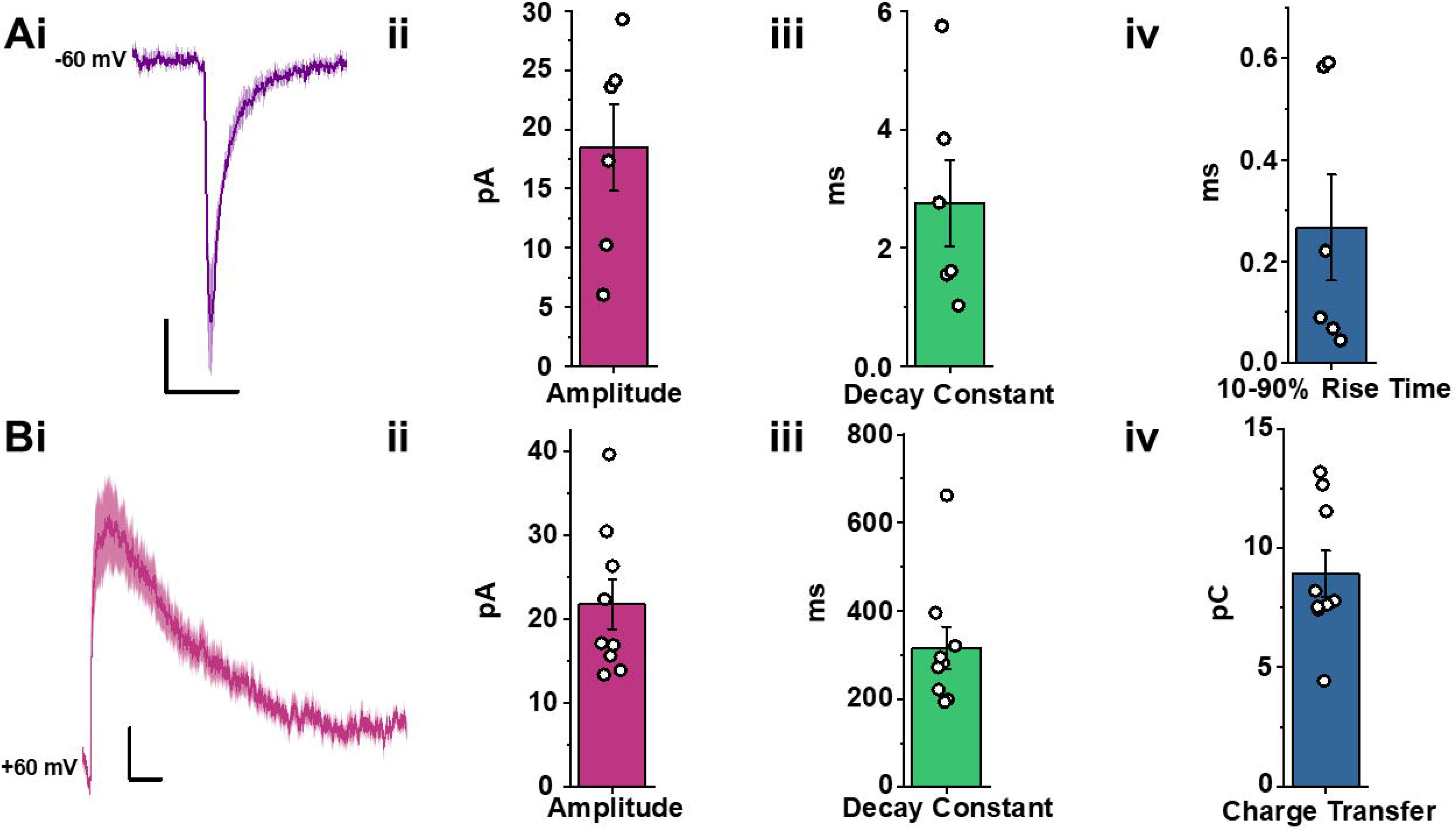
Whole-cell patch clamp recording of mEPSCs at -60mV and +60mV. A, B) Averaged mEPSCs of hSC superficial dorsal horn neurons recorded at -60mV (Ai, n = 6 cells from 4 male and 2 female donors) and at +60mV (Bi, n = 9 cells, with 7 cells from 5 male and 2 cells from 2 female donors). Scale bars = 10 ms (x axes), 5 pA (y axes). ii-iv) The amplitude, 10-90% rise time, and decay constant of mEPSCs measured from the corresponding traces.

The holding potential was then adjusted from −60 mV to +60 mV to record NMDAR-dominated mEPSCs. Figure 5Bi illustrates averaged mEPSCs (N = 9 cells) from 5 male and 2 female donors, displaying decay constants (315.61 ± 144.66 ms), charge transfer values (8.99 ± 2.90 pC), and amplitude measurements (21.72 ± 8.90 pA; N = 9) of saline-treated neurons (Figure 5Bii-iv). These experiments characterized the biophysical properties of NMDAR-mediated mEPSCs, which is important for understanding the kinetic profile and synaptic function of these currents in human SDH neurons. This foundational knowledge is critical for linking specific synaptic properties to functional outcomes.

#### Option B: hdMEA Recordings

High-density multi-electrode array (hdMEA) recordings are valuable for studying the electrophysiological properties of populations of neurons in an intact network, allowing for the high-resolution detection of neuronal activity and comprehensive analysis of spike properties across multiple sites. Here, we demonstrate the utility of hdMEAs for recording from hSC slices.

In Figure 6A, we show a quadrisected hSC slice mounted on the hdMEA chip, secured with a tissue anchor to ensure stable recordings. A representative spike trace from the SDH region is presented, allowing for the detailed assessment of spike amplitude and timing (Figure 6B).

**Figure 6.**
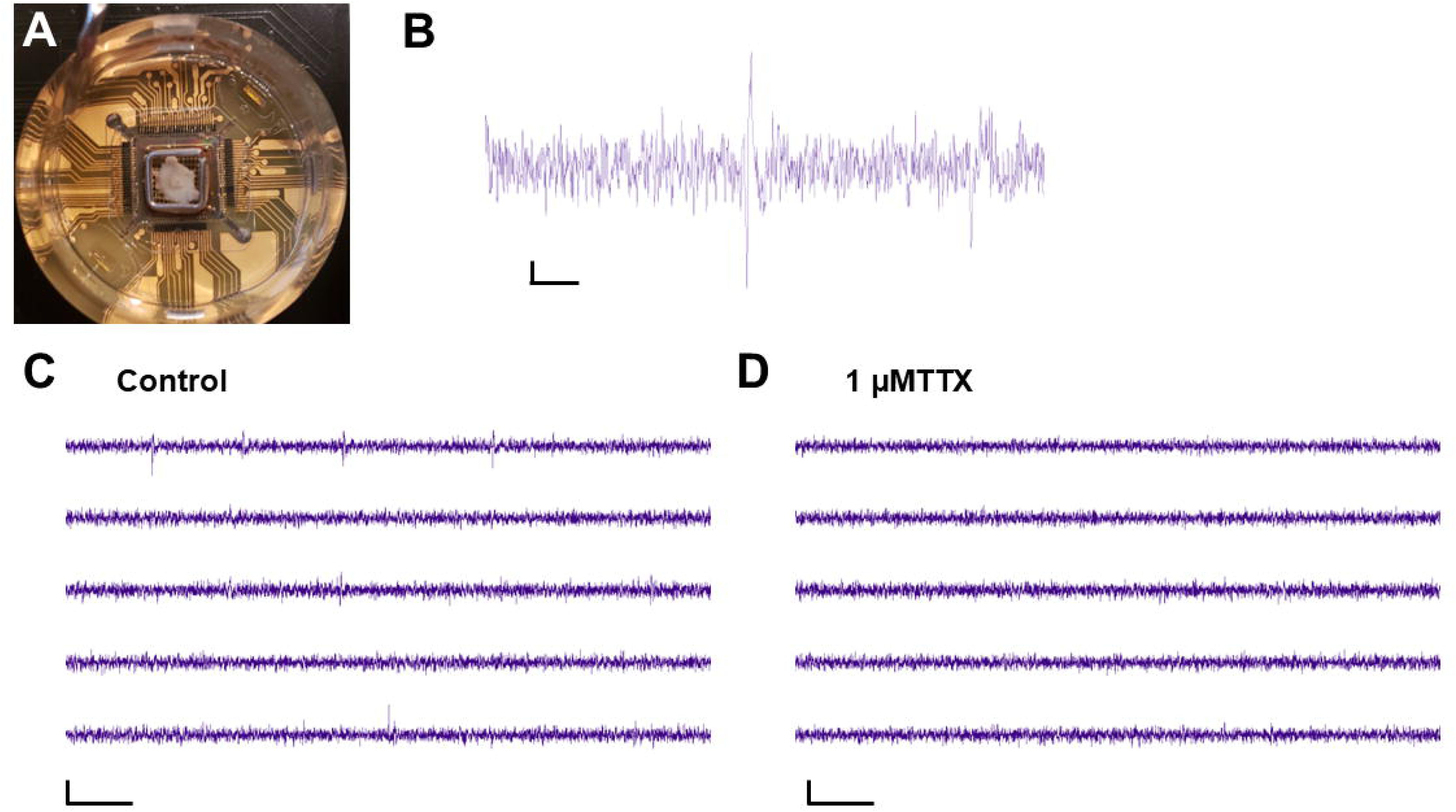
High-density multi electrode arrays (hdMEAs) can be used to investigate spike properties of hSC neurons. A) A hSC slice dorsal horn quadrisection mounted on the hdMEA chip, secured by a tissue anchor, in preparation for recording. B) A representative spike from the SDH region of a hSC slice. Scale bars: X axis =10ms, Y axis = 20µV. C) and D) Representative channels in the SDH region from a hSC recording following C) saline-treatment and following D) TTX-treatment. N = 1; Scale bars: X axis = 50 ms, Y axis = 100µV.

Figure 6C and D display representative activity from the hdMEA channels: saline-treated hSC tissue exhibited typical spiking activity (Figure 6C), while treatment with 1 μM tetrodotoxin (TTX) effectively abolished spiking, confirming that the recorded spikes represent neuronal action potentials (Figure 6D). The ability to capture individual spikes with hdMEAs enables precise characterization of spike properties and underlying neuronal excitability.

Using the hdMEA system for pharmacological experiments further demonstrates the versatility of hdMEA recordings for investigating dynamic changes in neuronal activity. We examined the effects of 2 µM capsaicin versus DMSO vehicle control, as well as the sodium channel blocker TTX, on the mean firing rate of SDH neurons in hSC slices (Figure 7A). Capsaicin increased neuronal firing (N = 5 recordings), while the DMSO vehicle did not (N = 2 recordings) (Figure 7B,C). TTX application completely inhibited activity, showcasing the time-course pharmacological responses over a 35-minute recording session (Figure 7C). The experiments, conducted using slices from two donors (one male, one female), illustrate the potential of hdMEAs for testing modulators like brain-derived neurotrophic factor (BDNF), which are known to drive hyperexcitability in male nociceptive networks of the hSC (Dedek et al., 2019, 2022).

**Figure 7.**
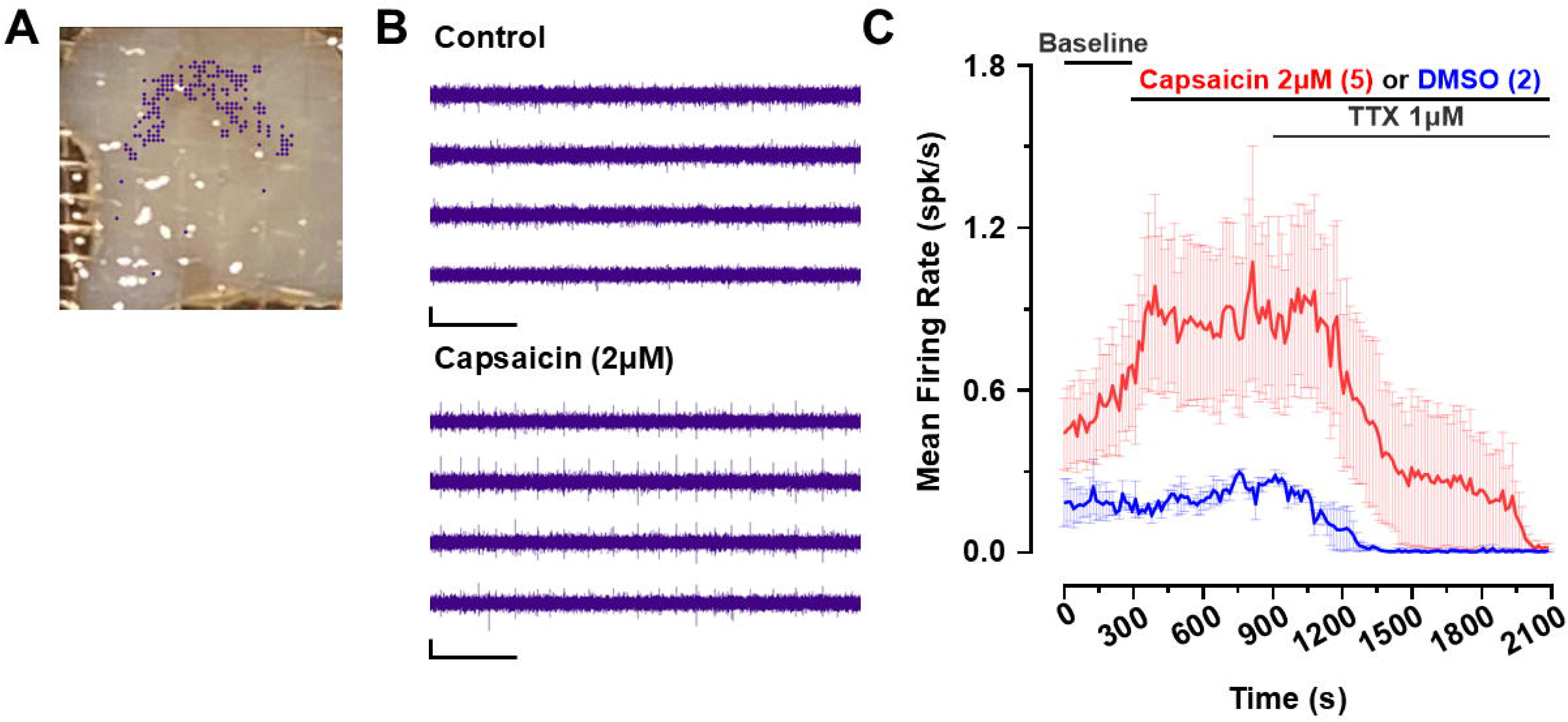
hdMEA recordings can be used for slice pharmacology experiments. A) hSC slice used for pharmacology experiments (data in C)) showing active channels in the SDH highlighted in purple. B) Sample traces from select active channels pictured in A) during the control (top) and after treatment with 2μM capsaicin (bottom). N =1 Scale bars: X axis = 1s, Y axis = 100µV. C) Pharmacological experiments showing the effects of 2μM capsaicin or DMSO vehicle and 1μM TTX on mean firing rate of hSC SPH neurons. Capsaicin N = 5 recordings, DMSO = 2 recordings; Recordings are from N = 2 donors (1 male, 1 female).

Overall, these tools provide a robust platform for exploring the impact of various agents on neuronal excitability. This capability is critical for identifying compounds that may inhibit pathological excitability, offering potential therapeutic candidates, or understanding mechanisms that exacerbate excitability, shedding light on pathways relevant to pain and other neurological disorders of the spinal cord.

## Author contributions

E.C.T. developed the surgical approach. A.D. and M.E.H optimized existing tissue preparation protocols for use in hSC. A.D., R.S., E.G., and M.E.H. collected data. E.T. helped analyze data. All authors contributed to writing and editing the manuscript and figures.

## Acknowledgments

We thank the organ and tissue donors and their families for their generous, selfless gift. We thank the Trillium Gift of Life Network, the surgical staff at The Ottawa Hospital Civic Campus and Dr. Suzan Chen, Lei Zhou, Dr. Ahmad Galuta, Dr. Sara Ameri, Jessica Parnell, and Dr.

Maitreya Patel for their help coordinating human spinal cord collection. Thank you to Christopher Dedek for his help in troubleshooting the human tissue slice preparation and patch-clamp electrophysiology.

