## Supplementary Figure 1 for "Processing and sectioning of organ donor spinal cord tissue for electrophysiology on acute human spinal cord slices" (from another version of this preprint: version 1, 2025-07-31)

**Supplementary Figures:**

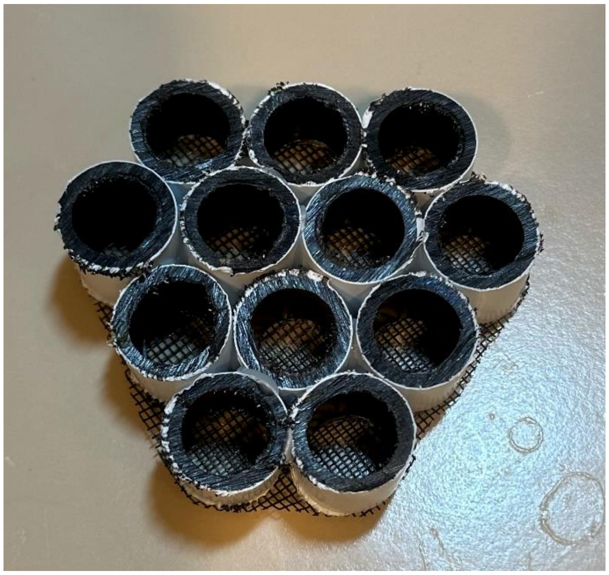

**Supplementary Figure 1.** Custom slice recovery chamber. This slice recovery chamber is made of 1” PVC piping glued onto a plastic screen. This size, with 12 individual chambers, fits into a 1L beaker. A smaller slice chamber is used to allow slices to passively to room temperature and contains 9 individual chambers.
